## Supplemental Table 1 and Sup Figs 1-4 for "Serial ‘deep-sampling’ PCR of fragmented DNA reveals the wide range of *Trypanosoma cruzi* burden among chronically infected hosts and allows accurate monitoring of parasite load following treatment"

S1 Table. Historical and year one PCR and hemoculture summary for macaques

| **ID** | **Sex** | **Birth Year** | **Year Infected/ Sero-converted** | **Age (yr) at infection** | **First confirmed conventional PCR pos.** | **Total number of PCR reactions (up to 12 months)** | **0verall % deep-sampling PCR pos** | **Ave positive Cq value** | **Overall % pos Hemoculture** |
| --- | --- | --- | --- | --- | --- | --- | --- | --- | --- |
| **P1** | M | 1999 | 2013 | 14 | 2016 | 60 | 70.00 | 31.92 | 54.3 |
| **P2** | M | 2000 | 2015 | 15 | 2016 | 3366 | 0.56 | 33.42 | 3.3 |
| **P3** | F | 2001 | 2013 | 12 | 2016 | 1,869 | 2.09 | 32.84 | 15.0 |
| **P4** | F | 2004 | 2014 | 10 | 2018 | 316 | 3.16 | * | * |
| **P5** | M | 2004 | 2015 | 11 | 2016 | 60 | 81.67 | 30.39 | 63.3 |
| **P6** | M | 2005 | 2012 | 7 | 2018 | 1784 | 5.60 | 33.71 | 1.7 |
| **P7** | F | 2005 | 2015 | 10 | 2018 | 3062 | 0.29 | 35.90 | 0 |
| **P8** | F | 2009 | 2014 | 5 | 2016 | 1980 | 1.31 | 34.07 | 5.7 |
| **P9** | M | 2009 | 2017 | 8 | Mult.neg. 2018 | 3256 | 0.21 | 33.44 | 1.7 |
| **P10** | F | 2011 | 2017 | 6 | 2018 | 712 | 4.77 | 32.66 | 15.0 |
| **P11** | M | 2012 | 2021 | 9 | no PCR data | 2888 | 1.76 | 35.03 | 15.0 |
| **P12** | M | 2013 | 2018 | 5 | no PCR data | 1592 | 0.25 | 31.20 | 0 |
| **P13** | M | 2013 | 2018 | 5 | no PCR data | 438 | 24.20 | 32.72 | 6.7 |
| **P14** | F | 2013 | 2021 | 8 | no PCR data | 70 | 52.86 | 29.70 | 73.3 |
| **P15** | F | 2014 | 2020 | 6 | no PCR data | 70 | 70.00 | 28.47 | 57.1 |
| **P16** | F | 2015 | 2021 | 6 | no PCR data | 70 | 81.42 | 25.59 | 48.6 |
| **P17** | M | 2015 | 2019 | 4 | no PCR data | 80 | 68.75 | 25.39 | 62.9 |
| **P18** | F | 2015 | 2019 | 4 | no PCR data | 60 | 56.70 | 32.11 | 46.7 |
| **P19** | F | 2015 | 2021 | 6 | no PCR data | 60 | 48.33 | 32.16 | 20.0 |
| **P20** | F | 2015 | 2021 | 6 | no PCR data | 70 | 40.00 | 32.64 | 43.3 |
| **P21** | F | 2015 | 2019 | 4 | no PCR data | 70 | 24.28 | 32.54 | 10.0 |
| **T1** | F | 2001 | 2011 | 10 | 2016 | 2310 | 0 |  | 0 |
| **T2** | F | 2005 | 2014 | 9 | 2016 | 1925 | 0 |  | 0 |
| **T3** | F | 2006 | 2015 | 9 | 2016 | 1925 | 0 |  | 0 |
| **T4** | F | 2006 | 2015 | 9 | 2016 | 1925 | 0 |  | 0 |
| **T5** | F | 2012 | 2016 | 4 | 2016 | 2310 | 0 |  | 0 |
| **N1** | F | 2009 | Never | NA | no PCR data | 2346 | 0.04 |  | 0 |
| **N2** | F | 2013 | Never | NA | no PCR data | 2343 | 0 |  | 0 |

*Died during study - chronic colitis, possible early colon cancer/precancerous lesion

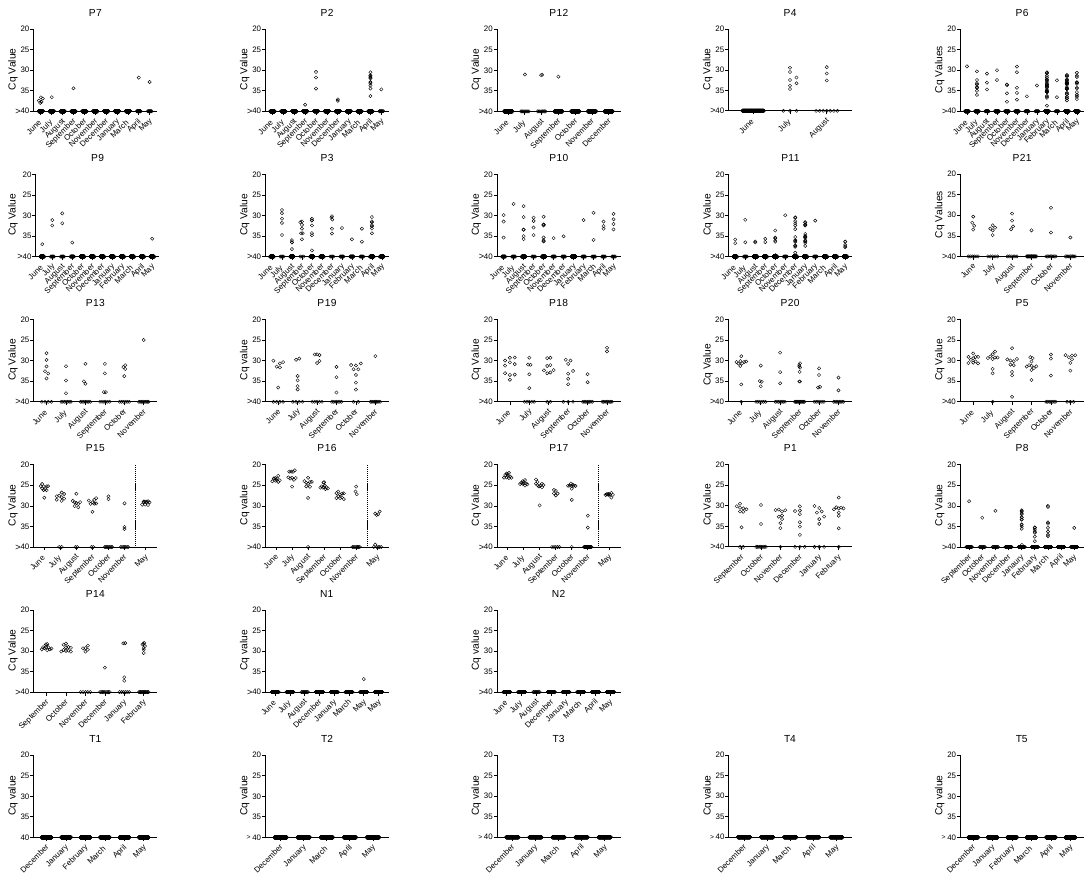

S1 Fig. The monthly pattern of detection of *T. cruzi* in replicate PCR reactions in DNA from macaque blood collected over one year of sampling. The positive reactions (Cq values <40) plotted for each month are the total from two blood samples collected each month (see Figure 2a). The total number of PCR reactions, the percent positive and the mean Cq values are provided in Sup table 1. N1 and N2 are seronegative controls and macaques T1-T5 were previously infected but cured of *T. cruzi* infection using benzoxaborole AN15368 [8]

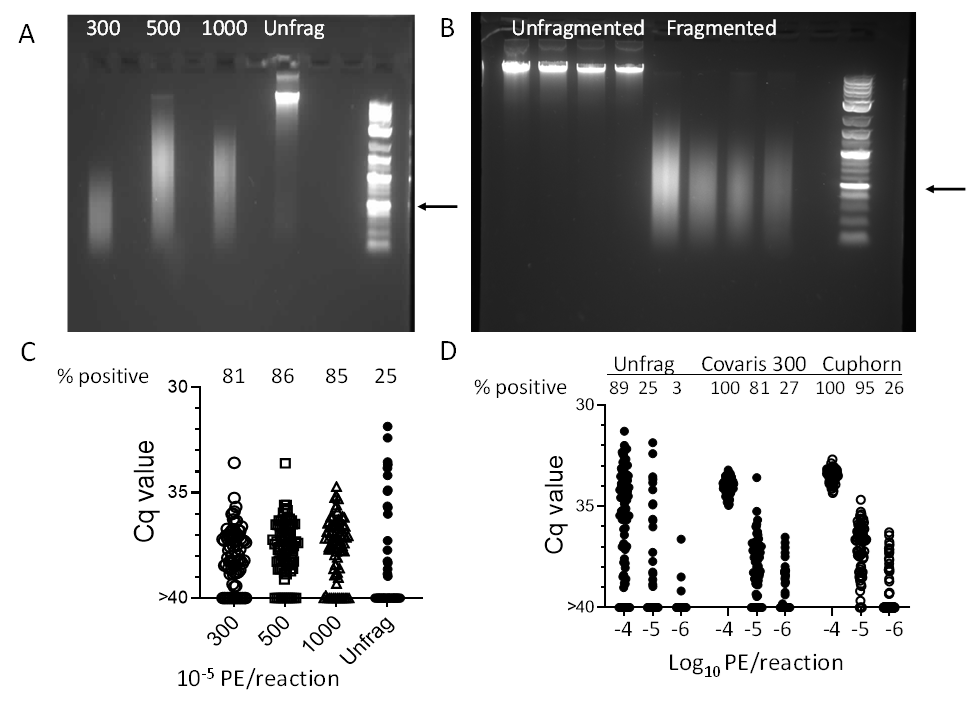

S2 Fig. Fragmentation of DNA increases the frequency of positive replicate PCR reactions in a DNA sample. A. Agarose gel profile of DNA isolated from macaque blood spiked with a known number of *T. cruzi* epimastigotes, either without fragmentation (Unfrag) or using Covaris settings to achieve an average fragment size of 300, 500 or 1000 bases. B. Four DNA samples from seropositive dogs pre- and post-fragmentation using a sonicator and cuphorn attachment as described in the Materials and Methods. Arrows indicate 500 bp band in DNA standards. C. Fragmentation at all three Covaris settings increased the frequency of PCR positive replicate aliquots with an average of 10^-5^ parasite equivalents per aliquot. A total of 78-81 replicates were amplified for each condition. D. The Covaris- and the cuphorn-fragmented DNA provided comparable increases in the frequency of product amplification at different parasite equivalents per aliquot as compared to unfragmented (Unfrag) DNA. A total of 72 to 90 aliquots were amplified for each condition.

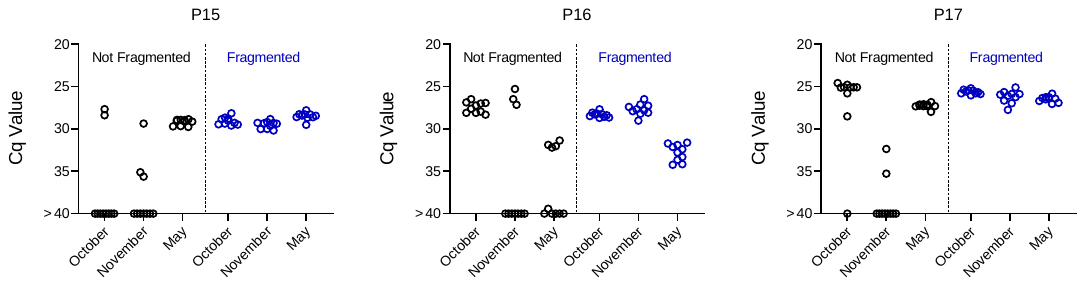

S3 Fig. DNA fragmentation results in more consistent PCR amplification and detection, even in samples where *T. cruzi* DNA may be detectable using a single PCR assay. Replicate aliquots (10 each) of non-fragmented and sonication-fragmented blood DNA from three macaques at three sampling times were amplified for detection of *T. cruzi* DNA.

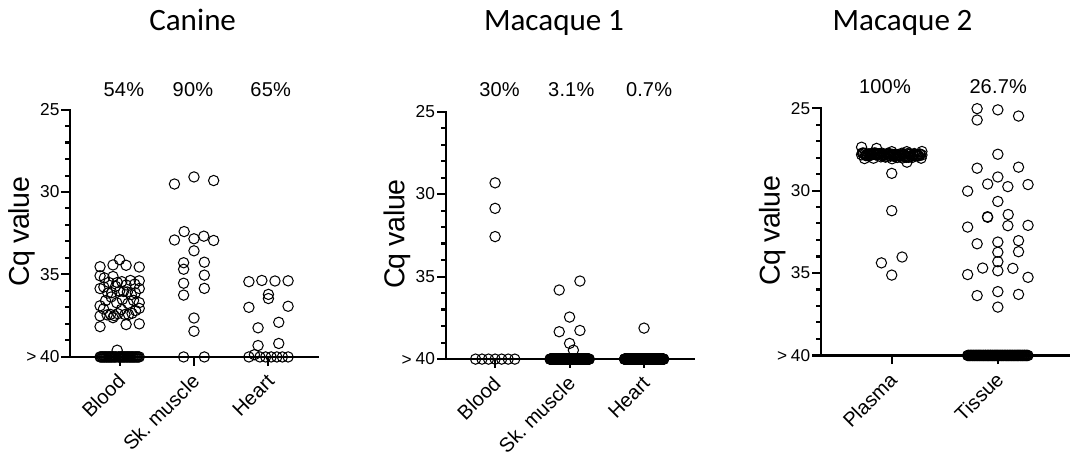

S4 Fig Detection of *T. cruzi* infection by deep-sampling PCR of blood- or plasma-derived DNA is corroborated by the PCR detection of *T. cruzi* DNA in individual tissue samples from skeletal muscle, heart, or other organs including liver, spleen and gut (Tissue). The indicated percentages reflect the fraction of PCR positive aliquots of a single DNA sample prepared from blood or plasma, or DNA purified from multiple individual sites in the case of muscle and tissues.
